# Targeted enhancement of the therapeutic window of L19-TNF by transient and selective inhibition of RIPK1-signaling cascade

**DOI:** 10.1101/688507

**Authors:** Sheila Dakhel, Tiziano Ongaro, Baptiste Gouyou, Mattia Matasci, Alessandra Villa, Dario Neri, Samuele Cazzamalli

**Author notes:** <u>Financial Support:</u> D.N. acknowledges funding from ETH Zurich. This project has received funding from the Swiss National Science Foundation (Grant Nr. 310030_182003/1) and the European Research Council (ERC) under the European Union’s Horizon 2020 research and innovation program (grant agreement 670603).

## Abstract

**Introduction:** Cytokine-based products are gaining importance for cancer immunotherapy. L19-TNF is a clinical-stage antibody-cytokine fusion protein that selectively accumulates to tumors and displays potent anticancer activity in preclinical models. Here, we describe an innovative approach to transiently inhibit off-target toxicity of L19-TNF, while maintaining antitumor activity.

**Methods:** GSK’963, a potent small molecule inhibitor of RIPK1, was tested in tumor-bearing mice for its ability to reduce acute toxicity associated with TNF signaling. The biological effects of L19-TNF on tumor cells, lymphocytes and tumor vessels were investigated with the aim to enable the administration of TNF doses, which would otherwise be lethal.

**Results:** Transient inhibition of RIPK1 allowed to increase the maximal tolerated dose of L19-TNF. The protective effect of GSK’963 did not affect the selective localization of the immunocytokine to tumors as evidenced by quantitative biodistribution analysis and allowed to reach high local TNF concentrations around tumor blood vessels, causing diffused vascular shutdown and hemorrhagic necrosis within the neoplastic mass.

**Conclusions:** The selective inhibition of RIPK1 with small molecule inhibitors can be used as a pharmaceutical tool to transiently mask TNF activity and improve the therapeutic window of TNF-based biopharmaceuticals. Similar approaches may be applicable to other pro-inflammatory cytokines.

## Introduction

The therapeutic benefit associated with PD-1 blockade in a growing number of cancer indications [1,2] has stimulated the search for alternative immunotherapy strategies, that may be complementary to the use of immune checkpoint inhibitors. Cytokines represent a group of small immunoregulatory proteins, some of which have been considered for cancer therapy applications [3]. Recombinant interleukin-2 has been shown to eradicate metastatic melanoma or renal cell carcinoma in a small proportion of patients, who are fit enough to tolerate the severe side-effects associated with cytokine treatment [4]. Recombinant tumor necrosis factor (TNF) in combination with melphalan has gained marketing authorization in Europe for the treatment of unresectable soft tissue sarcoma, in which TNF is administered through isolated limb perfusion procedure [5].

Recombinant cytokines may cause substantial toxicity at very low doses (sometimes less than 1 mg per patient), thus preventing escalation to therapeutically active regimens. For example, the maximal tolerated dose (MTD) of TNF for systemic administration procedures was found to be 300 μg/patient [6]. The fusion of cytokine payloads with recombinant antibodies, capable of selective localization to the tumor, may represent a strategy for the improvement of the therapeutic index of cytokine-based biopharmaceuticals [7,8]. Potent and selective anticancer activity of immunocytokine products has been observed in mouse models of cancer (particularly for fusion proteins based on IL2, IL12 and TNF payloads) [9–11]. Two fusion proteins of the L19 antibody (specific to the alternatively-spliced EDB domain of fibronectin, a pan-tumoral antigen whose expression in normal tissues is limited to female reproductive organs) are currently being investigated in Phase III clinical trials in Europe and in the United States. L19-IL2 has shown initial signs of activity against various tumor entities, including metastatic melanoma [12–14]. L19-TNF and other TNF-based fusion proteins were found to be potently active against soft-tissue sarcoma and the fully-human fusion protein is currently being studied in pivotal trials in this indication, in combination with doxorubicin (NCT03420014, Eudract no. 2016-003239-38).

The targeted delivery of TNF causes a rapid and selective hemorrhagic tumor necrosis in mouse models of cancer [15–17] and in patients [18]. TNF also boosts the activity of NK cells and of CD8+ T cells, which are required in order to eliminate the minimal residual disease and lead to protective immunity in mice [17]. However, TNF-based pharmaceuticals may cause toxicity, especially at early time points after intravenous administration, when the concentration in blood is highest. The most common side effects include a drop in blood pressure, flu-like symptoms, nausea and vomiting. These toxicities typically disappear when the blood concentration of the product falls below a critical threshold [19]. It would be desirable to develop strategies that preserve the therapeutic activity of targeted TNF pharmaceuticals, while decreasing systemic toxicity [7].

L19-TNF and related products provide a therapeutic benefit which is mediated by the durable high local concentration, that can be reached within the tumor mass. We reasoned that, if we were able to transiently inhibit TNF activity in blood (following intravenous administration of the product), we should be able to minimize side effects and safely administer higher doses of the product. One way to achieve this goal may be represented by the administration of small molecule inhibitors of key components in the TNF signaling pathway, building up activity at the tumor site, while decreasing systemic toxicity. Receptor-interacting protein kinase 1 (RIPK1) inhibitors may be ideally suited for this purpose, since RIPK1 is a key mediator of TNF-induced inflammation and tissue degeneration [20,21].

In this article, we report our findings on the combination of L19-hTNF (the clinical-stage fusion protein) or L19-mTNF (an analogue featuring murine TNF as therapeutic payload) with GSK’963, a selective RIPK1 inhibitor [22]. Combination therapy studies were also performed with ibuprofen, a non-steroidal anti-inflammatory drug, commonly used for the premedication of cancer patients undergoing cytokine treatment. The sequential administration of GSK’963 and L19-TNF did not inhibit the preferential accumulation of the antibody-cytokine fusion in the neoplastic mass and the induction of hemorrhagic necrosis in the tumor. The combination treatment allowed to safely administer L19-TNF beyond the maximal tolerated dose, with retention of potent anticancer activity.

By contrast, co-administration of L19-TNF and ibuprofen did not reduce body weight loss and treatment-related toxicity. The findings of this paper may be clinically relevant and may be applicable to other cytokine biopharmaceuticals. Combination regimens with judiciously-chosen small molecule inhibitors may allow to increase the maximal tolerated dose, thanks to a transient inhibition of signaling events in blood and in normal organs. A long residence time of the cytokine moiety within the tumor mass is less likely to be affected by the inhibitor, as small molecules are often cleared within minutes or hours.

## Material and Methods

### Compounds

GSK’963 (Aobious), GSK’2982772, Necrostatin-1 and Necrostatin-1s (Nec-1 and Nec-1s; Selleck Chemicals) were dissolve in 100% DMSO at 10 mg/ml and kept at −20°C for long-term storage. Ibuprofen sodium (Sigma-Aldrich) was prepared at 5 mg/ml in deionized water and stored at 4°C for no longer than 1 week.

### Protein Production

L19-human TNF (L19-hTNF) is a fusion protein consisting of the L19 antibody in scFv format, sequentially fused to human TNF by the 17-amino-acid linker EF(SSSSG)3 [23]. The antibody-cytokine fusion protein was produced as GMP material by Philogen S.p.A (Siena, Italy) and dialyzed with cellulose membranes (Spectra/Por^®^, Thermo Scientific) into phosphate-buffered saline (PBS; 20 mM NaH_2_PO_4_; 150 mM NaCl; pH 7.4) prior biodistribution experiments. L19-murine TNF (L19-mTNF) gene was cloned into the mammalian cell expression vector pcDNA3.1(+) (Invitrogen) by HindIII/NotI restriction sites. The L19-mTNF fusion protein was expressed by transient gene expression in CHO-S cells and purified from the cell culture supernatant to homogeneity by protein A (Sino Biological) chromatography, as described previously [24]. After dialyses into PBS pH 7.4, the quality of the proteins was assessed by SDS-PAGE, by Size-exclusion chromatography on a Superdex 200 Increase 10/300□LGL column on an ÄKTA FPLC (GE Healthcare) and by mass spectrometry (Waters Xevo G2XS Q-TOF) [**Supplementary Information**].

### Cell Lines

Chinese Hamster Ovary cells (CHO-S; Invitrogen) were cultured in suspension in PowerCHO-2CD medium (Lonza), supplemented with Ultraglutamine-1 (4 mM; Lonza) and antibiotic-antimycotic (1%; AA; Gibco).

Murine cells lines F9 (embryonal teratocarcinoma), WEHI-164 (fibrosarcoma) and CT-26 (colon carcinoma) were obtained from American Type Culture Collection. Cells were cultured in the corresponding medium supplemented with Fetal Bovine Serum (10%; FBS; Invitrogen) and AA (1%) following the supplier’s protocol and kept in culture for no longer than 10 passages with a confluence lower than 90%.

### *In Vitro* Cytotoxicity Assay

The direct killing activity of L19-mTNF was determined *in vitro* on the fibrosarcoma WEHI-164 cell line. Cells were seeded in 96-well plates (20’000 cells/well) in RPMI-164 medium (Gibco). After 24 hours, new medium containing decreasing concentration of L19-hTNF (1:10 dilution steps) with actinomycin D (Sigma; 2 μg/mL) was added to the cells, in the presence or absence of RIPK1 inhibitors (GSK’963, GSK’2982772, Nec-1 and Nec-1s; 1 μM). The cell viability was measured after 24 hours by adding 20 μl of CellTiter 96^®^ AQueous One Solution Reagent (Promega) directly to the wells. The number of living cells was determined by measuring the absorbance after 4 hours at 490 nm using Spark multimode microplate reader (Tecan). Experiments were performed in triplicate and the results were expressed as the percentage of cell viability compared with controls (cells treated with actinomycin D only).

### Animal Experiments

All the animal studies were conducted in accordance with Swiss animal welfare laws and regulations (license number ZH04/2018, granted by Veterinäramt des Kantons Zürich).

### Tumor Implantation

F9 cells were grown to 80% confluence and detached with Trypsin-EDTA 0.05% (Life Technologies). Cells were washed with Hank’s Balanced Salt Solution (HBSS, pH 7.4) once, counted and re-suspended in HBSS to a final concentration of 6.7 × 10^7^ cells ml^−1^. Aliquots of 10^7^ cells (150 μl of suspension) were injected subcutaneously (s.c.) in the right flank of female 129/Sv mice (6-7 weeks of age, Janvier).

WEHI-164 and CT26 cells were grown to 80% confluence and detached with Trypsin-EDTA 0.05% (Life Technologies). Cells were washed with Hank’s Balanced Salt Solution (HBSS, pH 7.4) once, counted and re-suspended in HBSS to a final concentration of 1.7 × 10^7^ cells ml^−1^. Aliquots of 2.5 × 10^6^ cells (150 μl of a suspension) were injected s.c. in the right flank of female BALB/c mice (6-7 weeks of age, Janvier).

Tumor volume was determined with the following formula: (d)^2^ × D × 0.52, where d and D are the short and long dimensions in millimeters, respectively, measured with a caliper. Animals were sacrificed when termination criteria foreseen by the license were reached.

### Cytokine analyses in plasma

Tumor-free BALB/c mice were injected intravenously (i.v.) with PBS, L19-mTNF (250 μg/Kg) or with GSK’963 (2 mg/Kg) followed by L19-mTNF (250 μg/Kg; 30 min later). After 2 hours, mice were euthanized and immediately exsanguinated. Blood was collected and incubated for 20 min in Microtainer tubes containing lithium heparin (BD Microtainer Tube). Plasma was obtained by centrifugation for 15 min at 3’000 rpm using a refrigerated centrifuge and stored at −80°C prior to cytokine quantification.

IL2, IL6, IL10, MCP-1, IFN-γ TNF and IL1β cytokine plasma levels were quantified using a multiplexed bead-based flow cytometric assay kit (Bio-Techne AG, Zug, Switzerland) following the described procedure [25] at the Cytolab facility (Regensdorf, Switzerland).

### Quantitative Biodistribution Experiment

The *in vivo* targeting performance of L19-hTNF was evaluated in biodistribution studies as previously described [26]. Briefly, the immunocytokine was radio-iodinated with ^125^I (Hartmann Analytic) and Chloramine T hydrate (0.25 μg/μg protein, Sigma) and purified on a PD10 column (GE Healthcare). Radiolabeled L19-hTNF (250 μg/Kg) was injected into the lateral tail vein of immunocompetent 129/Sv mice bearing s.c. implanted F9 teratocarcinoma. Alternatively, mice were pretreated with the RIPK1 inhibitor GSK’963 (2 mg/Kg) followed by i.v. injection of L19-hTNF (250 μg/Kg; 30 min later). After 24 hours mice were sacrificed, organs were excised, weighed and the radioactivity of organs and tumors was counted using a Packard Cobra gamma counter. Radioactivity content of representative organs was expressed as the percentage of the injected dose per gram of tissue (%ID/g).

### Therapy Experiments

WEHI-164 tumor-bearing BALB/c mice were randomized in groups of four and received three i.v. injections, once every second day. Mice were pretreated with either GSK’963 (2 mg/Kg) or ibuprofen at (5 mg/Kg) 30 min prior to injection of L19-mTNF. Recommended (250 μg/Kg) or high (375 μg/Kg) doses of L19-mTNF were tested. Toxicity induced by murine TNF was monitored daily by checking the body weight of the animals. A body weight loss of more than 10% was considered as severe consequence of the treatment. Tumors were measured with an electronic caliper to assess antitumor efficacy of the different treatments. Daily statistics are described in the **Supplementary Information**.

### Immunofluorescence Studies

For the *ex vivo* detection of vascular permeability, WEHI-164 or CT-26 bearing BALB/c mice were injected i.v. with PBS, L19-mTNF (250 μg/Kg) or with GSK’963 (2 mg/Kg) followed by L19-mTNF (250 μg/Kg; 30 min later). After 24 hours, 10 mg/Kg of Hoechst 33342 (Thermo Scientific) was injected into the tail vein, and animals were euthanized after 1 min. Tumor, kidney and liver were collected, embedded in cryoembedding medium (Thermo Scientific) and snap frozen in liquid nitrogen. Cryo-sections of 10□m were fixed in ice-cold acetone for 10 min and blocked for 1 hour in FBS (20% solution in PBS). The vessels were stained with a rat anti-mouse CD31 (BD Pharmigen; 1:100) and with a donkey anti-rat Alexa Fluor 594 as a secondary antibody (Invitrogen; 1:500). Slides were mounted with fluorescent mounting medium (Dako, Glostrup, Denmark) and analyzed with Leica TIRF epi-fluorescence microscope (Scientific Center for Optical and Electron Microscopy ScopeM, ETH, Switzerland). Pictures representing a sample overview were obtained by stitching electronically adjacent regions.

### Statistics

Data were analyzed using Prism 7.0 (GraphPad Software, Inc.). Student t test was used to assess the differences of cell viability and of cytokine levels between different experimental groups. In therapy experiments, statistical significances were determined with a regular 2-way ANOVA test (with Bonferroni post-test; *, p≤0.05; **, p≤0.01; ***, p≤0.001; and ****, p≤0.0001).

## Results

### Production of L19-TNF fusion proteins and quality control

L19-hTNF and L19-mTNF were produced in CHO cells and purified to homogeneity [**Figure 1**]. Sequences of the fusion proteins are described in the **Supplementary Information**. The proteins, which form stable non-covalent homotrimers thanks to the TNF moiety, were pure in SDS-PAGE analysis and eluted as single peak in gel-filtration. Deconvolution of LC-MS profile allowed identification of the expected experimental masses for L19-mTNF and L19-hTNF, which were compatible with the predicted values in the absence of any protein glycosylation [**Figure 1D** and **1E**].

**Figure 1:**
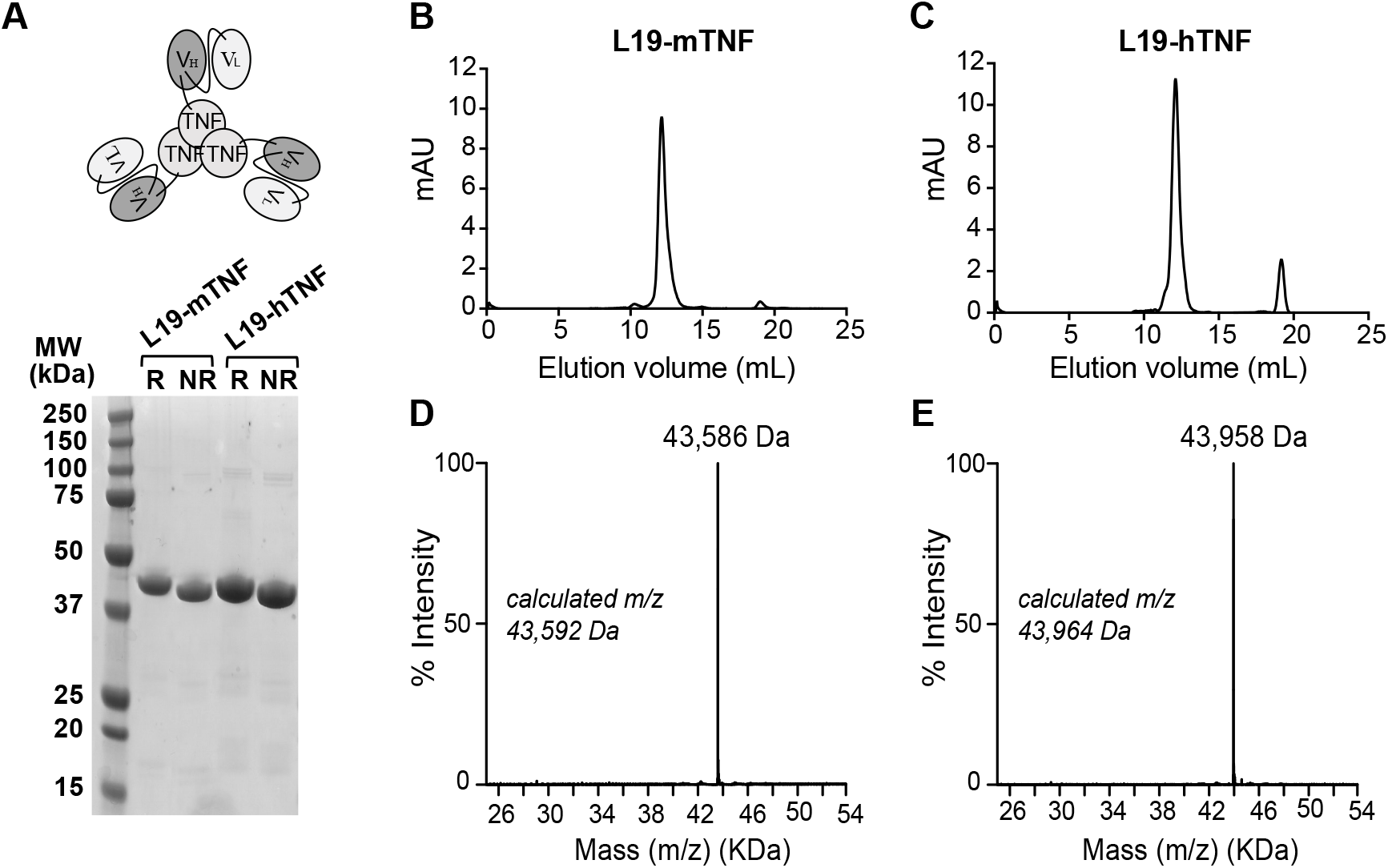
Quality control analyses of L19-TNF fusion proteins. (A) Schematic representation of L19-TNF and SDS-PAGE analysis. (B, C) Size exclusion chromatography and (D, E) ESI-MS profile of L19-mTNF and of L19-hTNF, respectively. MW, molecular weight; R, reducing conditions; NR, nonreducing conditions.

### RIPK1 inhibitors reduce *in vitro* potency of L19-TNF

Small molecule inhibitors of RIPK1 [22,27,28], a key kinase in the signaling cascade of TNF through its TNF receptor 1 (TNFR1) [**Figure 2A**], were tested *in vitro* for their ability to reduce potency of L19-mTNF. Cytotoxicity assays were performed on the murine fibrosarcoma WEHI-164 cell line. All four tested inhibitors (GSK’963, GSK’2982772, Nec-1 and Nec-1s) potently reduced TNF-mediated biocidal activity in a dose-dependent manner [**Figure 2B**]. Inhibition of the biocidal TNF activity by RIPK1 small molecule inhibitors was confirmed also for L19-hTNF on the WEHI-164 cell line [**Supplementary Figure 1**]. For further investigations, we chose GSK’963 as combination partner for L19-TNF, since this molecule was slightly more active than the other inhibitors and since extensive information (e.g., metabolic stability, pharmacokinetic properties, efficacy *in vivo)* was available for this molecule [22].

**Figure 2:**
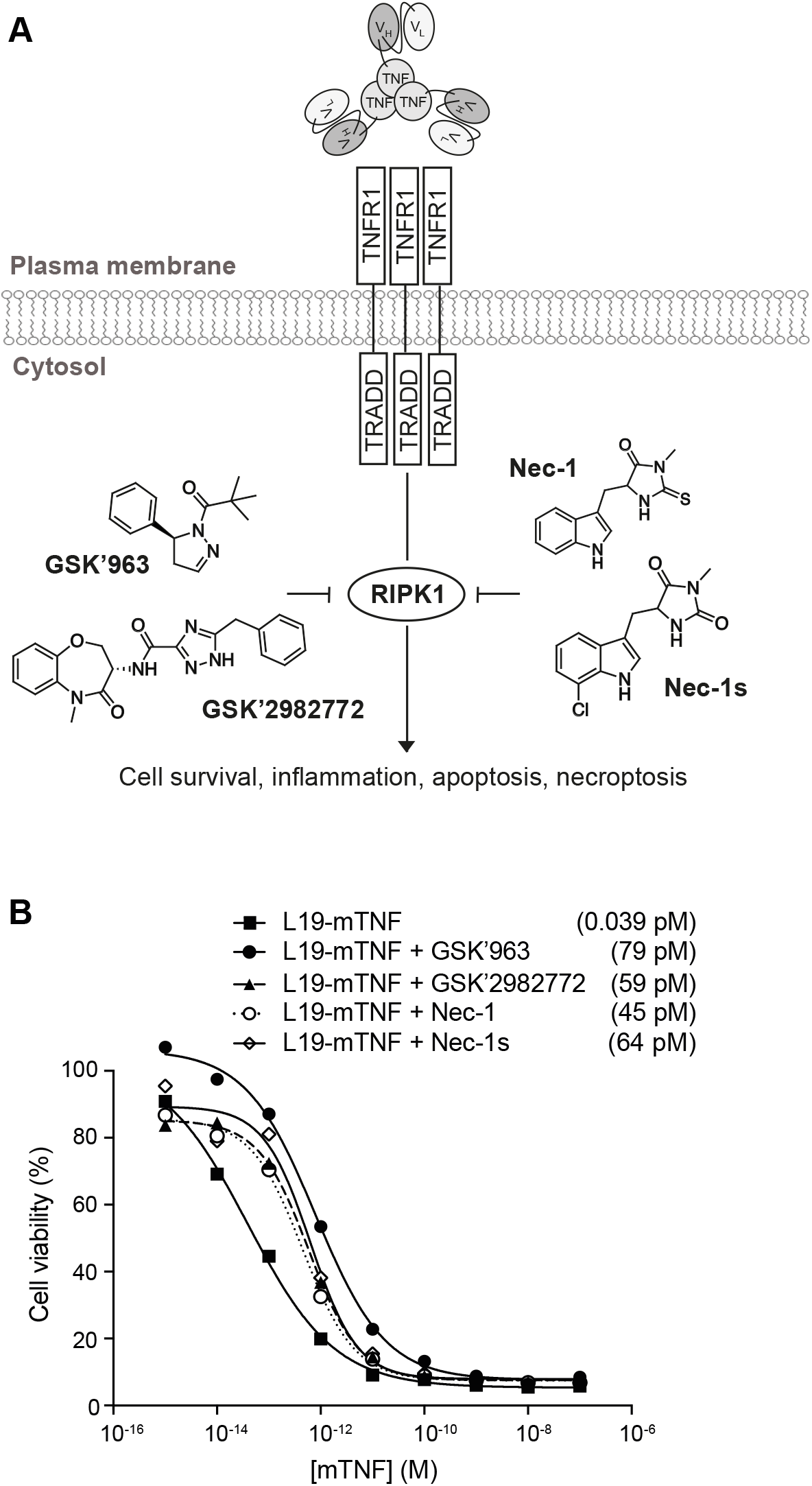
*In vitro* biocidal effect of L19-TNF. (A) Schematic representation of L19-TNF and TNFR1. The interaction between TNF and its receptor triggers a cascade of intracellular events which can be blocked by small molecule inhibitors of RIPK1 (structures of common RIPK1 inhibitors considered in this article are depicted). (B) *In vitro* activity of L19-mTNF alone or in combination with small molecule RIPK1 inhibitors. Dose-response curves of L19-mTNF (□□) on WEHI-164 murine fibrosarcoma obtained in the presence or absence of 1μM of GSK’963 (•), GSK’2982772 (▲), Necrostatin-1 (○) or Necrostatin-1s (Ͷ). Each data value represents the mean of cell viability ± SD (n=3). In all cases, tested inhibitors of RIPK1 were able to reduce the killing activity of targeted-TNF. The potency of L19-mTNF is expressed as calculated IC50 value in brackets.

### GSK’963 does not inhibit the ability of L19-mTNF to induce pro-inflammatory cytokines production *in vivo*

To characterize the immunological response to intravenously injected L19-mTNF, we measured plasma levels for a panel of relevant cytokines in immunocompetent tumor-free mice. L19-mTNF (given at the recommended dose of 250 μg/Kg) potently induced high levels of pro-inflammatory cytokines (e.g., IL6, MCP-1 and TNF) already two hours after the injection, compared with the non-treated control group [**Figure 3A**]. Pre-treatment with GSK’963 did not change the cytokine level profile, compared with the L19-mTNF monotherapy group. However, MCP-1 (monocyte chemoattractant protein 1) slightly decreased when animals were treated with combinatorial treatment of L19-mTNF and GSK’963 (p<0.01).

**Figure 3:**
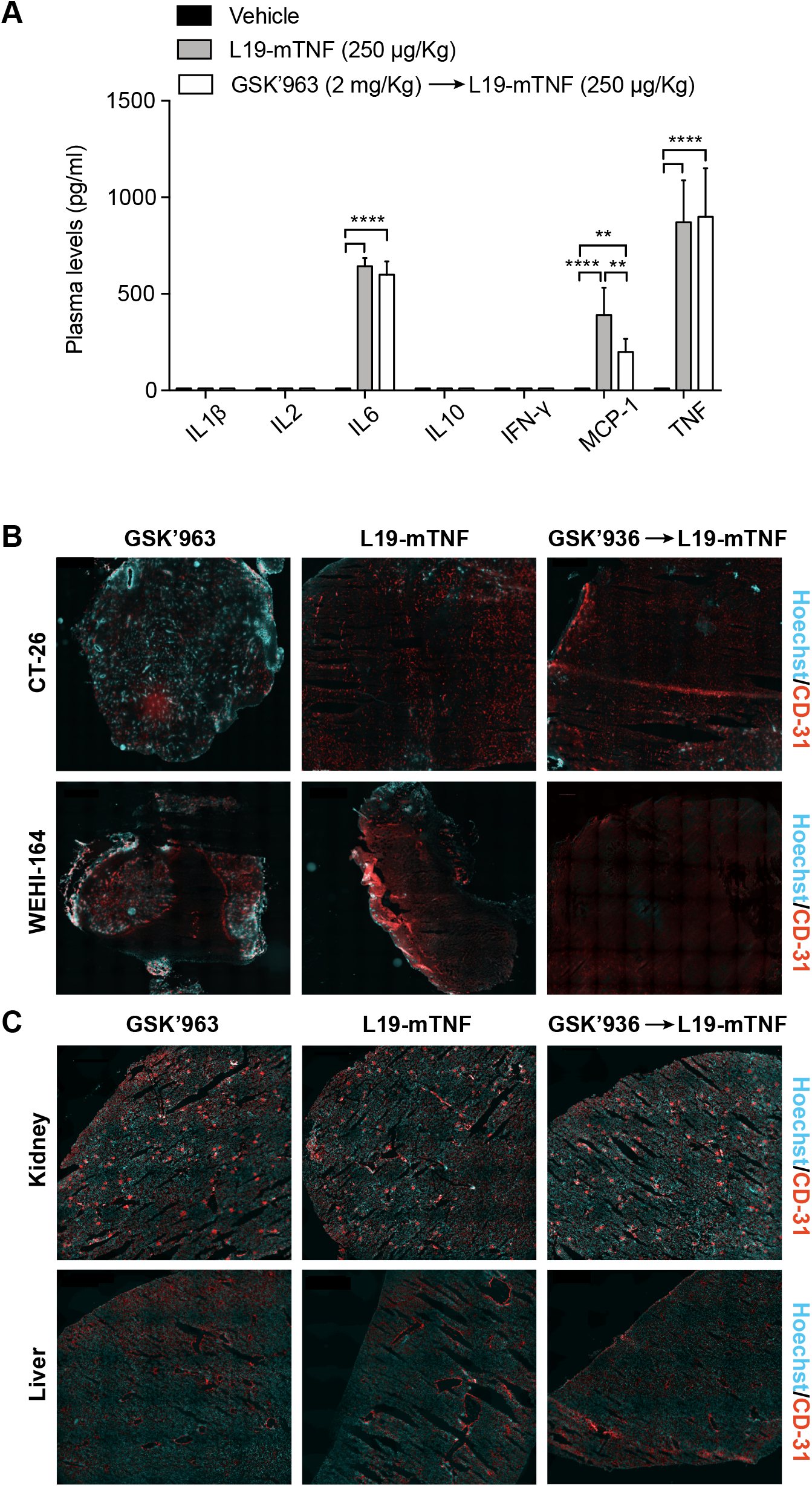
Determination of L19-mTNF effects on cytokine levels and vasculature in mice. The immunocytokine was given alone or in combination with the small molecule GSK’963, a potent small molecule inhibitor of RIPK1. (A) Quantification of plasma cytokine concentrations in immunocompetent BALB/c mice 2 hours after treatment with L19-mTNF. Data represent mean concentration values (± SD; n = 4 mice per group). Statistical differences were evaluated between groups (****, p≤0.0001; **, p≤0.01). (B) Vascular permeability study in tumors (CT-26 and WEHI-164), healthy kidney and liver. Hoechst was injected i.v. 1 min before the sacrifice to analyze the permeability of tumor and organ vasculature. Pictures represent an overview obtained by stitching electronically adjacent regions of the samples (20x magnification; scale bars 500 μm). Blue = Hoechst staining; Red = CD31 vessels staining.

### GSK’963 does not interfere with tumor-specific vascular shut-down induced by L19-TNF

To gain more insight on the effects of L19-mTNF on tumor and normal blood vessels, we collected healthy organs and tumor samples (CT-26 and WEHI-164) 24 hours after *in vivo* administration of the immunocytokine as single agent or in combination with GSK’963. Hoechst 33342 dye was perfused one minute prior to sacrifice, in order to assess variations in the perfusion and functionality of blood vessels. Vascular structures were detected by CD31 staining. Administration of L19-mTNF at the recommended dose of 250 μg/Kg (alone or combined with GSK’963) prevented penetration of the Hoechst dye in both CT-26 and WEHI-164 tumors, indicating the onset of a selective vascular shutdown in neoplastic lesions [**Figure 3B**]. By contrast, no differences in vascular permeability were observed in kidney and liver between the different treatment groups [**Figure 3C**]. Apoptotic cell death was detected in tumor and healthy organs (kidney and liver) by immunofluorescence staining of Caspase-3 after the different treatments. Tumors treated either with L19-mTNF alone or in combination with GSK’963 were characterized by high number of dead cells (Caspase-3 positive in green), in contrast with neoplastic samples excised from animals in the untreated group (PBS). Apoptosis was not detectable in healthy organs following L19-mTNF administration [**Supplementary Figure 2**].

### Pre-treatment with GSK’963 is compatible with *in vivo* selective tumor accumulation of L19-TNF

The tumor-targeting performance of L19-TNF in combination with GSK’963 was evaluated in immunocompetent 129/Sv mice bearing subcutaneously-grafted F9 tumors. L19-hTNF was radiolabeled with ^125^I and injected intravenously at the recommended dose of 250 μg/Kg. L19-hTNF preferentially localized at the site of the disease, with a high tumor uptake value (18% of the injected dose/gram of tissue; %ID/g) and excellent tumor-to-normal organs ratio, 24 hours postadministration (i.e. average tumor-to-organs ratio 5.5:1 and tumor-to-blood ratio 4:1) [**Figure 4**]. Pre-treatment with GSK’963 (2 mg/Kg; i.v. 30 min prior systemic administration of the immunocytokine) did not substantially alter the distribution profile of L19-TNF.

**Figure 4:**
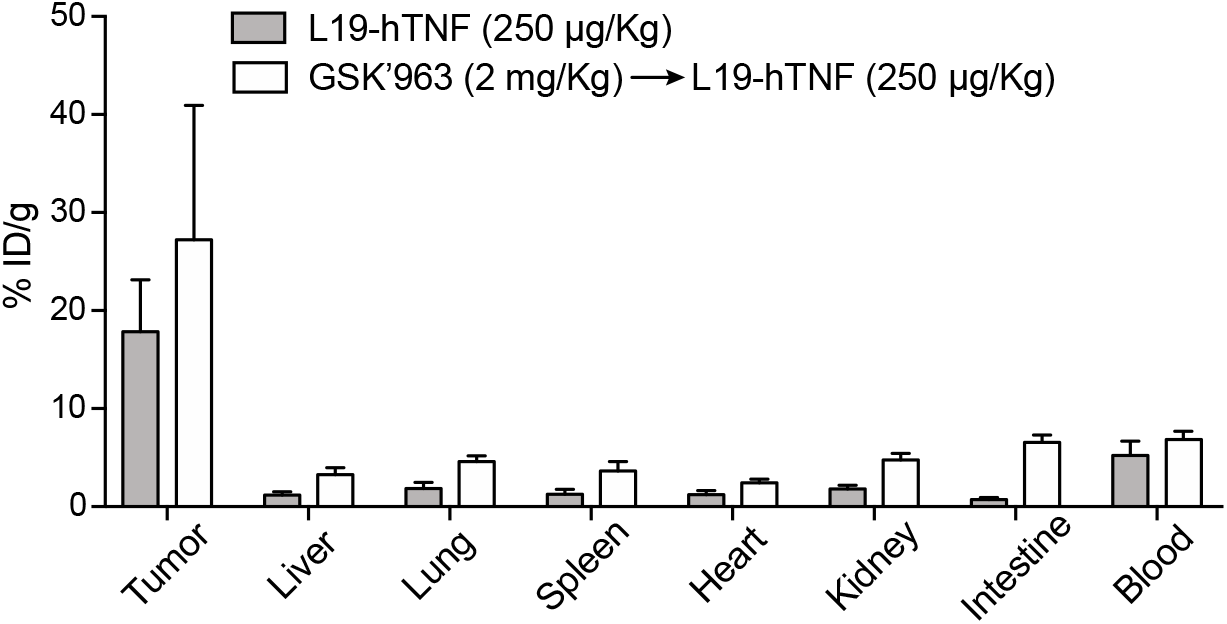
Biodistribution profile of targeted-TNF in F9 tumor-bearing mice. The immunocytokine was administered as monotherapy or in combination with GSK’963. Drug uptake values in tumors and healthy organs are expressed as percentage of the injected dose per gram of tissue (average; %ID/g ± SD) measured 24 hours post-injection.

### The therapeutic window of L19-TNF is improved by GSK’963

In order to determine the maximum tolerated dose of L19-TNF in combination with GSK’963, we performed a therapy study in WEHI-164 tumor-bearing mice. Animals were dosed systemically with L19-mTNF at 250 and 375 μg/Kg, respectively. Two additional groups of animals were treated with GSK’963 (2 mg/Kg) 30 min before administration of L19-mTNF. Acute toxicity was evaluated by measuring daily changes in body weight. Administration of L19-mTNF at the recommended dose (250 μg/Kg) caused reversible body weight loss (<10%, corresponding to moderate toxicity) [**Figure 5A**], while the high dose treatment (L19-mTNF at 375 μg/Kg) led to severe toxicity (body weight loss >15%) already 24 hours after the first administration [**Figure 5B**]. Pre-administration of GSK’963 resulted in complete protection from TNF-induced body weight loss at the recommended dose of L19-mTNF [**Figure 5A**]. Moreover, the severe acute toxicity induced by high dose of L19-mTNF was masked by pre-treatment with GSK’963, with increased overall survival of the animals [**Figure 5B**]. The tumor volumes were measured along experiment in order to compare the anticancer activity of the different treatments. The use of a combinatorial setting with GSK’963 [**Figure 5C** and **5D**] did not interfere with the therapeutic potential of the immunocytokine compared to monotherapy. In a parallel therapy experiment, we replaced pre-treatment with GSK’963 by administration of the general anti-inflammatory drug ibuprofen (5 mg/Kg), 30 min prior to L19-mTNF injection (375 μg/Kg). In this case, ibuprofen was not able to revert the severe body weight loss [**Figure 5E** and **5F**] induced by administration of a high dose of L19-mTNF.

**Figure 5:**
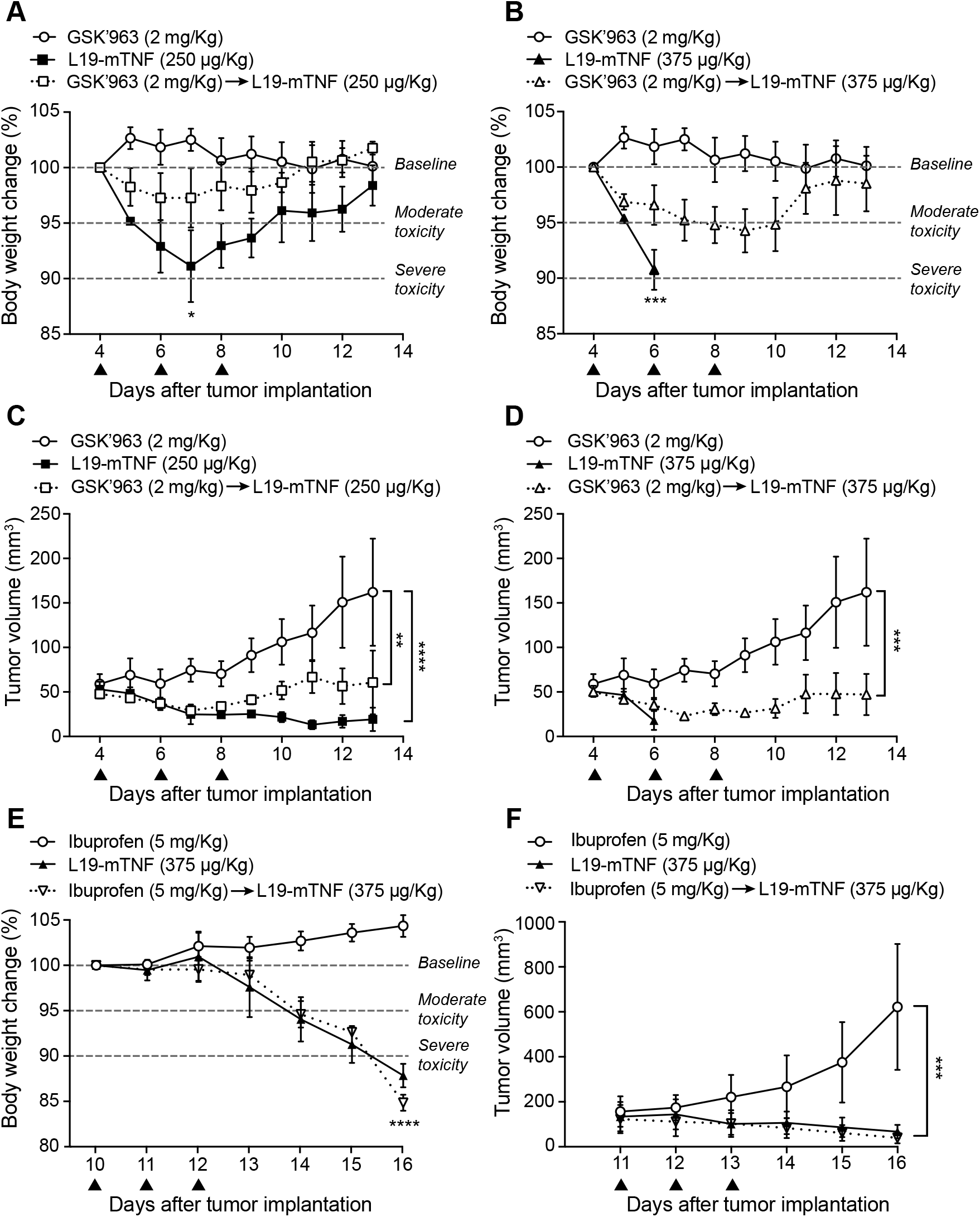
Tolerability and efficacy study in a syngeneic subcutaneous model of WEHI-164 fibrosarcoma. Different doses of L19-mTNF were given i.v. alone or in combination with GSK’963 (▲ indicates i.v. administration of corresponding compounds). (A, B) Changes in animal body weight induced by treatments at the recommended (250 *μ*/Kg) and at a high (375 *μ*/Kg) dose of L19-mTNF, respectively. (C, D) Plot of tumor volume measured daily and corresponding to different treatment groups. The same *in vivo* model was used to compare the L19-mTNF at high dose as a monotherapy or in a combinatorial setting with ibuprofen. Body weight changes (E) and tumor volume (F) were analyzed daily. Data represent mean of the experimental values (± SEM; n = 4 mice per group). Statistical differences were assessed between treatment and control groups (****, p≤0.0001; ***, p≤0.001; **, p≤0.01; *, p≤0.05).

## Discussion

We have described the use of a potent small molecule inhibitor of RIPK1 named GSK’963 for the improvement of the therapeutic window of an antibody-TNF fusion protein (L19-TNF) in preclinical models of cancer. Combination with GSK’963 enabled dose escalation of the immunocytokine beyond its MTD, while the single-dose administration of an equivalent amount of L19-TNF (375 *μ*g/Kg) was found to be lethal. Inhibition of RIPK1 did not interfere with selective accumulation of L19-TNF into neoplastic lesions, where high local concentrations of TNF caused onset of tumor vascular shutdown and selective hemorrhagic necrosis. In contrast with these findings, pre-administration of ibuprofen at its recommended dose did not have a similar beneficial effect in reducing toxicities related to the administration of targeted-TNF.

TNF is a potent cytokine mediator of inflammation which is expressed by macrophages, NK cells, T cells, endothelial cells and fibroblasts [29]. Trimeric TNF interacts with two different receptors, TNFR1 and TNFR2, causing activation of complex signaling pathways that overall account for TNF-induced cell death, inflammation and cell activation. Tissue degeneration and inflammation is mainly promoted by the interaction of TNF with TNFR1 and by subsequent recruitment and activation of the intra-cellular RIPK1 protein [30].

The use of recombinant TNF has been previously proposed for cancer therapy, but significant toxicities and lack of efficacy prevented its clinical success for systemic application [31]. Hypotension, leukopenia, thrombocytopenia, fever, headache, nausea and hepatopathy represent the most common side effects detected in Phase II studies with recombinant TNF, while most serious toxicities include respiratory failure and coagulopathies [32]. Our group has previously demonstrated that the generation of antibody-TNF fusion proteins capable to selectively accumulate to tumors after systemic administration represents a valuable strategy to improve therapeutic window and efficacy of TNF [15–17]. L19-TNF is an antibody-fusion protein for the targeting of the alternatively-spliced EDB domain of fibronectin [33], that shows potent antitumor activity in preclinical models of cancer [9,34,35] and is now being investigated in clinical trials [13,18,36]. The product entered Phase III clinical trials in combination with L19-IL2 as neoadjuvant therapy prior to surgery of fully-resectable Stage IIIB,C melanoma (NCT02938299 and NCT03567889). Moreover, the combination of L19-TNF plus doxorubicin is being compared to doxorubicin monotherapy in pivotal trials as first-line treatment of advanced soft-tissue sarcoma (EudraCT 2016-003239-38 and NCT03420014).

While therapeutic activity can be dramatically improved by the selective delivery of cytokines to tumors via antibodies [15–17], toxicity profile is often similar to the one of unmodified cytokines. The majority of side effects of immunocytokines are typically observed in correspondence with the peak serum concentration of the product [13,36]. Transient symptoms including chills, fever, fatigue and pain are for example observed already few minutes after L19-TNF infusion in patients, with a peak around 1-2 hours after application and a total duration of few hours [36]. Pharmacological inhibition of cytokine activity shortly after intravenous administration represents an opportunity to improve the therapeutic window. Small molecules (like GSK’963) are particularly attractive for this application, since their serum half-life matches the clearance rate of recombinant antibody products from circulation [37] and inhibition of the cytokine activity may be harmonized with and prevent the onset of systemic side-effects of immunocytokines [22,28,38].

We have successfully demonstrated that the MTD of L19-TNF can be increased by transient and selective inhibition of RIPK1, a key mediator of TNF-induced inflammation and tissue damage. Moreover, our results clearly show that small molecule inhibitors of RIPK1 are more effective in limiting the early toxicity of L19-TNF, as compared to general anti-inflammatory drugs like ibuprofen. Our results may be of clinical relevance, as patients treated with L19-TNF may benefit from judicious combinations with RIPK1 inhibitors, with potential improvement in terms of efficacy and safety of the treatment. Indeed, pharmacological approaches similar to the ones described in this article for L19-TNF may be applicable to other TNF-based pharmaceuticals or to other cytokine-based products. In particular, toxicities related to the administration of IL12-or IL2-antibody products [14,24,39,40] might benefit from the transient and selective inhibition of key intracellular mediators of their activity.

## Acknowledgments

The authors wish to thank Charlotte Howell for technical support in the production of L19-mTNF. The authors acknowledge support of the Scientific Center for Optical and Electron Microscopy ScopeM of the Swiss Federal Institute of Technology ETHZ. We personally acknowledge Justine Kusch for the help with microscopy. D.N. acknowledges funding from ETH Zurich, from the Swiss National Science Foundation (Grant Nr. 310030_182003/1) and from the European Research Council (ERC) under the European Union’s Horizon 2020 research and innovation program (grant agreement 670603).

