## Supplementary figure 1 and 2 for "Targeted enhancement of the therapeutic window of L19-TNF by transient and selective inhibition of RIPK1-signaling cascade"

**SUPPLEMENTARY INFORMATION**

Sheila Dakhel^a^, Tiziano Ongaro^a^, Baptiste Gouyou^a^, Mattia Matasci^a^, Alessandra Villa^a^, Dario Neri^b^, Samuele Cazzamalli^a^*

a) Philochem AG, Libernstrasse 3, CH-8112 Otelfingen (Switzerland)

b) Department of Applied Biosciences, Swiss Federal Institute of Technology (ETH Zürich), Vladimir-Prelog-Weg 4, CH-8093 Zurich (Switzerland)

*) Corresponding author

### Abbreviations

| AA | Antibiotic Antimycotic |
| --- | --- |
| Da  DMSO | Daltons  Dimethylsulfoxide |
| ESI-ToF-MS  FA  FCS | Electrospray ionization time-of-flight Mass Spectrometry  Formic Acid  Fetal Calf Serum |
| HBSS  I | Hank’s Balanced Salt Solution  Iodine |
| IC  LC-MS | Inhibitory Capacity  Liquid Chromatography-Mass Spectrometry |
| MeCN  MeOH | Acetonitrile  Methanol |
| MPa  MS | Mega Pascal  Mass Spectroscopy |
| MTS  MW | 3-(4,5-dimethylthiazol-2-yl)-5-(3-carboxymethylphenyl)-2-(4-sulfophenyl)-2H-tetrazolium)  Molecular Weight |
| OD  PBS | Optical Density  Phosphate-Buffered Saline |
| RPMI | Roswell Park Memorial Institute |
| s.c.  scFv  SD  SDS  SEM | Subcutaneous  Single Chain Fragment Variable  Standard Deviation  Sodium Dodecyl Sulfate  Standard Error of the Mean |
| TNFR  UPLC | Tumor Necrosis Factor Receptor  Ultra-Performance Liquid Chromatography |

### Protein Production and Purification

The L19-mTNF fusion protein (sequence reported below) was cloned into the mammalian expression vector pcDNA3.1(+) (Invitrogen) using a strategy similar to the one described before [1]. The protein was expressed in CHO-S cells (Invitrogen) by transient gene expression. Briefly, CHO cells in suspension were first counted and resuspended in fresh ProCHO medium to a final cell concentration of 4 ✕ 10^6^ cells/mL. 0.75 μg DNA/million cell and 2.5 μg PEI/million cells were added carefully to the cells. Cells were incubated in a shaker at 31 °C ✕ 150 rpm for 6 days. After incubation the suspension was centrifuged at 4 °C ✕ 4000 rpm for 30 minutes (JA-10 rotor) using AVANTI J-26S XP centrifuge (Beckman Coulter). Supernatant was filtered with 0.45μM filters (Nalgene) and incubated for 2h at room temperature with Protein A agarose beads resin (Sino Biological) before loading onto the PD-10 column. The column was thereafter washed with 200 mL of Buffer A (100 mM NaCl, 0.5 mM EDTA, 0.1% Tween 20 in PBS) and then with 200 mL Buffer B (500 mM NaCl, 0.5 mM EDTA in PBS). The antibody product was eluted using 10-15 mL 0.1 M glycine at pH = 3 and fractions of 1 mL were collected. OD at an absorbance of 280 nm (OD_280_) was measured and fractions containing protein (OD_280_ > 0.1 mg/mL) were pooled and loaded on SpectraPor dialysis membrane MW 3500 (Spectrum laboratories) and dialyzed in PBS o/n at 4 °C. After dialysis, L19-mTNF was characterized by SDS-PAGE, size exclusion chromatography, surface plasmon resonance and mass spectrometry.

### Protein Sequence of L19-mTNF

EVQLLESGGGLVQPGGSLRLSCAASGFTFSSFSMSWVRQAPGKGLEWVSSISGSSGTTYYADSVKGRFTISRDNSKNTLYLQMNSLRAEDTAVYYCAKPFPYFDYWGQGTLVTVSSGDGSSGGSGGASEIVLTQSPGTLSLSPGERATLSCRASQSVSSSFLAWYQQKPGQAPRLLIYYASSRATGIPDRFSGSGSGTDFTLTISRLEPEDFAVYYCQQTGRIPPTFGQGTKVEIKSSSSGSSSSGSSSSGLRSSSQNSSDKPVAHVVANHQVEEQLEWLSQRANALLANGMDLKDNQLVVPADGLYLVYSQVLFKGQGCPDYVLLTHTVSRFAISYQEKVNLLSAVKSPCPKDTPEGAELKPWYEPIYLGGVFQLEKGDQLSAEVNLPKYLDFAESGQVYFGVIAL

### Protein Sequence of L19-hTNF

EVQLLESGGGLVQPGGSLRLSCAASGFTFSSFSMSWVRQAPGKGLEWVSSISGSSGTTYYADSVKGRFTISRDNSKNTLYLQMNSLRAEDTAVYYCAKPFPYFDYWGQGTLVTVSSGDGSSGGSGGASEIVLTQSPGTLSLSPGERATLSCRASQSVSSSFLAWYQQKPGQAPRLLIYYASSRATGIPDRFSGSGSGTDFTLTISRLEPEDFAVYYCQQTGRIPPTFGQGTKVEIKEFSSSSGSSSSGSSSSGVRSSSRTPSDKPVAHVVANPQAEGQLQWLNRRANALLANGVELRDNQLVVPSEGLYLIYSQVLFKGQGCPSTHVLLTHTISRIAVSYQTKVNLLSAIKSPCQRETPEGAEAKPWYEPIYLGGVFQLEKGDRLSAEINRPDYLDFAESGQVYFGIIAL

### Protein Characterization

#### SDS-PAGE

Protein samples were diluted to 0.2-0.3 mg/mL in PBS and mixed with either reducing or non-reducing 5x Loading buffer. Samples were denatured 5’ at 95 °C and loaded on NuPAGE 4-12% Bis-Tris Gel (Novex™ by Life Technologies). 1x MES NuPAGE (Novex™ by Life Technologies) was used as running buffer and electrophoresis was performed at 180 V, 110 mA for 1h. Gel was rinsed with deionized water and stained in Coomassie blue for 15-20’ on an orbital shaker. Staining solution was discarded and the gel was rinsed 3 times with deionized water and immerged in destaining solution (10% acetic acid/30% methanol/mQ water) for 3-12h on an orbital shaker. Destaining solution was discarded and recycled, gel was rinsed with deionized water and a picture of the gel was taken. Recipes for the 5X Loading buffer and Coomassie blue stain are as described in Table S1 and S2:

| **100mL, 5X non-red Loading Buffer** | |
| --- | --- |
| Tris­HCl (250mM, pH 6.8) | 20.8 mL |
| Glycerol | 33.3 mL |
| SDS | 6.6 g |
| Bromophenol blue | 66 mg |
| mQ water | up to 100 mL |

Table S1. 5X Loading buffer recipe. For 5X reducing Loading buffer, add 5-10% (v/v) 2-­mercaptoethanol

| **1L coomassie blue** | |
| --- | --- |
| PlusOne Coomassie PhastGel Blue R-350 | 2 tablets |
| Methanol | 400 mL |
| Acetic Acid | 100 mL |
| mQ water | 500 mL |

Table S2. Coomassie blue recipe

#### Gel Filtration Analysis

100 μL of diluted sample (final concentration 0.1-0.5 mg/mL) were loaded on FPLC (Äkta, GE Healthcare) and protein were separated by a Superdex200 Increase 10/300 GL column (GE Healthcare) previously equilibrated with 1 CV PBS, using PBS as mobile phase at a flow rate of 0.6 mL/min (column pressure limit set at 5 MPa). Proteins were detected by an UV-detector at a wavelength of 280 nm.

#### Mass Spectrometry

Samples were diluted to about 0.1 mg/mL and LC-MS was performed on a Waters Xevo G2XS Qtof instrument (ESI-ToF-MS) coupled to a Waters Acquity UPLC H-Class System using a 2.1 × 50 mm Acquity BEH300 C4 1.7 µm column (Waters). 0.1% FA in water (solvent A) and 0.1% FA in MeCN (solvent B) were used as mobile phase at a flow rate of 0.4 mL/min. Gradient was programmed as follows: after 1.5 min isocratic with 95% solvent A, stepwise change from 95% solvent A to 95% solvent B in 4.5 min (10% increase every 0.5 min), back to 95% solvent A in 0.5 min, linearly to 95% solvent B and back to 95% solvent A in 2.25 min (last step repeated twice).

### *In vitro* cytotoxicity assays of L19-hTNF on WEHI-164 in combination with RIPK1 inhibitors

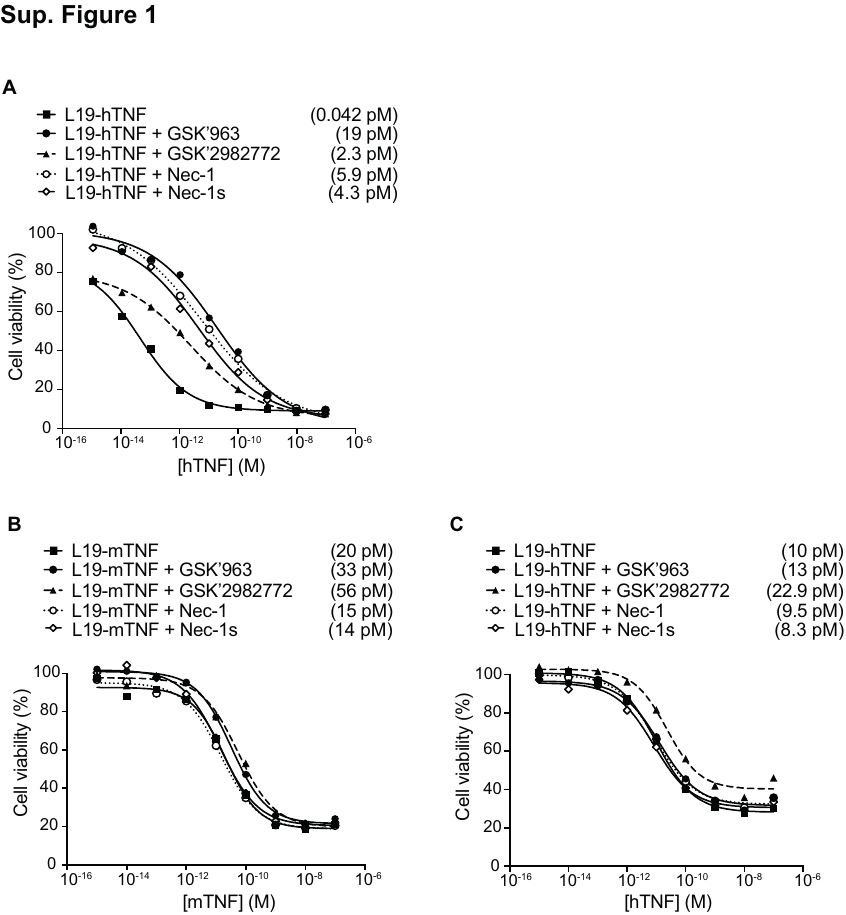

**Supplementary Figure 1: *In vitro* activity of L19-hTNF alone or in combination with small molecule RIPK1 inhibitors.** Dose-response curves of L19-hTNF (◼︎) obtained in the presence or absence of 1µM of GSK’963 (●), GSK’2982772 (▲), Necrostatin-1 (○) or Necrostatin-1s (◇) on WEHI-164 murine fibrosarcoma. Each data value represents the mean of cell viability ± SD (n = 3). The potency of L19-hTNF is expressed as calculated IC50 values in brackets.

### *Ex vivo* immunofluorescence apoptosis analysis

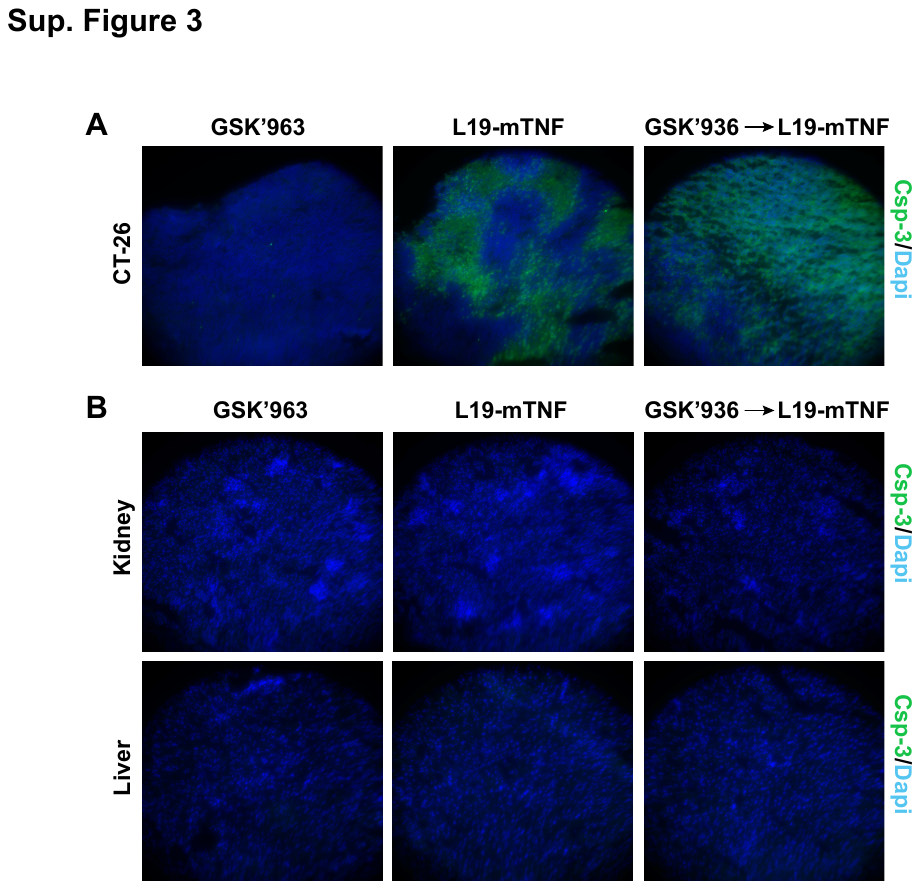

**Supplementary Figure 2** Apoptosis staining after L19-mTNF treatment (250 µg/Kg) alone or in combination with GSK’963 (2 mg/Kg) in CT-26 tumor-bearing mice (n = 1 per group). Twenty- four hours after the i.v. injections, healthy organs (kidney and liver) and tumors were excised and processed as describe before in the material and methods. Slides were stained using a rabbit anti-caspase-3 antibody (apoptotic cells; 1:200 dilution; Sigma) and detected with a goat anti-rabbit AlexaFluor488 secondary antibody (green; 1:500 dilution; Invitrogen). Nuclei were counterstained with DAPI (blue; 1:1000; Invitrogen). Pictures represent a significant region of the entire sample (20x magnification).

### Statistical analysis of therapy experiments

Differences in tumor volume and body weight between therapeutic groups were compared using the two-way ANOVA analysis with Bonferroni post-test of Graphpad Prism 7 (La Jolla, CA, USA). Days are counted after tumor implantation.

**Tumor Size (mg)**

GSK’963 (2 mg/Kg) vs L19-mTNF (250 µg/Kg).

From day 4 to day 9	non-significant differences

day 10	p < 0.05

day 11	p < 0.001

from day 12	p < 0.0001

GSK’963 (2 mg/Kg) vs GSK’963 (2 mg/Kg) 🡪 L19-mTNF (250 µg/Kg).

From day 4 to day 11	non-significant differences

From day 12	p < 0.01

L19-mTNF (250 µg/Kg) vs GSK’963 (2 mg/Kg) 🡪 L19-mTNF (250 µg/Kg).

From day 4 to day 13	non-significant differences

GSK’963 (2 mg/Kg) vs L19-mTNF (375 µg/Kg).

day 4	non-significant differences

day 5	non-significant differences

day 6	p < 0.05

GSK’963 (2 mg/Kg) vs GSK’963 (2 mg/Kg) 🡪 L19-mTNF (375 µg/Kg).

From day 4 to day 9	non-significant differences

day 10	p < 0.05

day 11	non-significant differences

from day 12	p < 0.001

L19-mTNF (375 µg/Kg) vs GSK’963 (2 mg/Kg) 🡪 L19-mTNF (375 µg/Kg).

From day 4 to day 6	non-significant differences

L19-mTNF (250 µg/Kg) vs L19-mTNF (375 µg/Kg).

From day 5 to day 6	non-significant differences

L19-mTNF (250 µg/Kg) vs GSK’963 (2 mg/Kg) 🡪 L19-mTNF (375 µg/Kg).

From day 4 to day 13	non-significant differences

GSK’963 (2 mg/Kg) 🡪 L19-mTNF (250 µg/Kg) vs GSK’963 (2 mg/Kg) 🡪 L19-mTNF (375 µg/Kg).

From day 4 to day 13	non-significant differences

**Body Weight Change (%)**

GSK’963 (2 mg/Kg) vs L19-mTNF (250 µg/Kg).

day 4	non-significant differences

day 5	p < 0.05

day 6	p < 0.01

day 7	p < 0.0001

day 8	p < 0.01

day 9	p < 0.01

From day 10	non-significant differences

GSK’963 (2 mg/Kg) vs GSK’963 (2 mg/Kg) 🡪 L19-mTNF (250 µg/Kg).

From day 4 to day 13	non-significant differences

L19-mTNF (250 µg/Kg) vs GSK’963 (2 mg/Kg) 🡪 L19-mTNF (250 µg/Kg).

From day 4 to day 13	non-significant differences

GSK’963 (2 mg/Kg) vs L19-mTNF (375 µg/Kg).

day 4	non-significant differences

day 5	p < 0.001

day 6	p < 0.0001

GSK’963 (2 mg/Kg) vs GSK’963 (2 mg/Kg) 🡪 L19-mTNF (375 µg/Kg).

From day 4 to day 6	non-significant differences

day 7	p < 0.05

day 8	non-significant differences

day 9	p < 0.05

from day 10	non-significant differences

L19-mTNF (375 µg/Kg) vs GSK’963 (2 mg/Kg) 🡪 L19-mTNF (375 µg/Kg).

day 4	non-significant differences

day 5	non-significant differences

day 6	p < 0.05

L19-mTNF (250 µg/Kg) vs L19-mTNF (375 µg/Kg).

From day 4 to day 6	non-significant differences

L19-mTNF (250 µg/Kg) vs GSK’963 (2 mg/Kg) 🡪 L19-mTNF (375 µg/Kg).

From day 4 to day 13	non-significant differences

GSK’963 (2 mg/Kg) 🡪 L19-mTNF (250 µg/Kg) vs GSK’963 (2 mg/Kg) 🡪 L19-mTNF (375 µg/Kg).

From day 4 to day 13	non-significant differences

**Tumor Size (mg)**

Ibuprofen (5 mg/Kg) vs L19-mTNF (375 µg/Kg).

From day 10 to day 14	non-significant differences

day 16	p < 0.001

Ibuprofen (5 mg/Kg) vs ibuprofen (5 mg/Kg) 🡪 L19-mTNF (375 µg/Kg).

From day 10 to day 15	non-significant differences

day 16	p < 0.001

L19-mTNF (375 µg/Kg) vs ibuprofen (5 mg/Kg) 🡪 L19-mTNF (375 µg/Kg).

From day 10 to day 16	non-significant differences

**Body Weight Change (%)**

Ibuprofen (5 mg/Kg) vs L19-mTNF (375 µg/Kg).

From day 10 to day 13	non-significant differences

day 14	p < 0.001

from day 15	p < 0.0001

Ibuprofen (5 mg/Kg) vs ibuprofen (5 mg/Kg) 🡪 L19-mTNF (375 µg/Kg).

From day 10 to day 13	non-significant differences

day 14	p < 0.001

from day 15	p < 0.0001

L19-mTNF (375 µg/Kg) vs ibuprofen (5 mg/Kg) 🡪 L19-mTNF (375 µg/Kg).

From day 10 to day 16	non-significant differences
